# Consumption of carotenoids not increased by bacterial infection in brown trout embryos (*Salmo trutta*)

**DOI:** 10.1101/293142

**Authors:** Lucas Marques da Cunha, Laetitia G. E. Wilkins, Laure Menin, Daniel Ortiz, Véronique Vocat-Mottier, Claus Wedekind

## Abstract

Carotenoids are organic pigment molecules that play important roles in signalling, control of oxidative stress, and immunity. Fish allocate carotenoids to their eggs, which gives them the typical yellow to red colouration and supports their resistance against microbial infections. However, it is still unclear whether carotenoids act mainly as a shield against infection or are used up during the embryos’ immune defence. We investigated this question with experimental families produced from wild-caught brown trout (*Salmo trutta*). Singly raised embryos were either exposed to the bacterial pathogen *Pseudomonas fluorescens* or sham-treated at one of two stages during their development. A previous study on these experimental families reported positive effects of egg carotenoids on embryo growth and resistance against the infection. Here, we quantified carotenoid consumption in these infected and sham-infected maternal sib groups. We found that carotenoid contents mostly decreased during embryogenesis. However, these decreases were neither linked to the virulence induced by the pathogen nor dependent on the time point of infection. We conclude that egg carotenoids are not significantly used up by the embryos’ immune defence.

## 1 Introduction

Carotenoids are naturally occurring pigments that have been shown to be relevant for a wide range of physiological functions in animals. They are involved in (i) antioxidant activity by facilitating singlet oxygen quenching and free radicals scavenging [1, 2], (ii) immune system functioning by, for example, protecting immune cells and maintaining an efficient immune response [3, 4] and (iii) retinol biosynthesis, which plays an important role in the immune system, vision, and embryo development [5, 6]. Many fishes, including most salmonids, allocate carotenoids to their eggs [7, 8]. For example, in Chinook salmon (*Oncorhynchus tshawytscha*), females mobilize carotenoids from their muscles to their eggs a few weeks before the spawning season, which produces the typical yellow to red egg coloration [9]. Egg carotenoids may therefore be an important component of maternal environmental effects in fish [5]. However, their role during embryo development is not sufficiently understood yet.

Salmonids are excellent models to study maternal effects. Females produce large numbers of eggs that are externally fertilized. Therefore, (i) experimental studies based on *in vitro* fertilizations are possible, (ii) embryos can be raised in the laboratory where potentially confounding factors such as female differential investment after zygote formation can be controlled for [10], (iii) embryos can be singly raised at high replication, which enables powerful statistical analyses [e.g. 11], and (iv) natural populations can be sampled, which allows studying the variation in egg contents within their ecologically relevant range. Previous studies found strong parental effects on embryo performance in response to different types of environmental stressors, with maternal effects typically being larger than paternal effects [e.g. 12, 13, 14]. The relative relevance of maternal effects decreases throughout embryogenesis, possibly due to the depletion of maternally-allocated compounds to the eggs and increasing relevance of paternal, i.e., additive genetic effects [15].

Experimental studies on captive populations and based on supplementary feeding of carotenoids have found positive links between egg carotenoids and offspring performance [16–19]. However, there seem to be non-linear dose dependencies, i.e., high amounts of some carotenoids can be equally or even less beneficial than intermediate levels [20–22]. This suggests that supplementary feeding could potentially produce artefacts. Wilkins et al. [23] and Wilkins et al. [24] have therefore sampled natural populations of brown trout (*Salmo trutta*) and studied the variance of naturally allocated egg carotenoids. They found astaxanthin, lutein, and zeaxanthin to be present in the eggs of all females. Capsanthin was found only in the eggs of few individuals. Importantly, egg carotenoid contents varied significantly among females (some females allocated several times more of certain carotenoids to the eggs than others). Wilkins et al. [23] and Wilkins et al. [24] investigated the potential significance of this variation in carotenoids for embryo performance. In their first study [23], embryos were stressed by an experimentally induced organic pollution. Embryo mortality was high and positively correlated to loss in carotenoids during embryogenesis (as determined at the level of the family and on the surviving embryos), but there was no correlation between embryo mortality and the initial carotenoid content in the eggs. The causalities of the links between carotenoids and stress-induced mortality remained unclear because the consumption of carotenoids in embryos that had died remained unknown [23]. Their second study on a new sample of eggs [24] used a low-virulence strain of *Pseudomonas fluorescens* as a stressor to avoid the constraints produced by high mortalities. This second study also measured offspring traits beyond embryo survival. *P. fluorescens* is an opportunistic pathogen that naturally occurs on brown trout eggs [25] and that can be used in experimental infections [11, 15]. Wilkins et al. [24] found astaxanthin egg contents to be positively correlated to larval growth in all treatment groups. Moreover, astaxanthin contents seemed to protect embryos from the virulence caused by the pathogen. *P. fluorescens* induced a delay in hatching time that was negatively correlated to the egg content of this carotenoid. However, it remained unclear whether astaxanthin just prevented stress (i.e., reduced susceptibility to the pathogen) or whether it was metabolized in the defense mechanism against the infection [5].

If carotenoids only prevented stress, we would expect a positive correlation between carotenoid contents in unfertilized eggs and indicators of stress resistance but no link between carotenoid consumption and stress resistance. If, however, carotenoids were used up in the response to stress, we would expect carotenoid consumption to be correlated to the stress resistance. A positive correlation between indicators of stress resistance and carotenoid consumption would indicate that carotenoids were used up in the response to stress but also that carotenoid availability was often limited. A negative correlation between stress resistance and carotenoid consumption would imply that embryos varied in their primary susceptibility to the stress. This variation in susceptibility might be due to embryo genetics [15] or other maternal environmental effects, such as egg size [26] or other compounds that females had allocated to their eggs before spawning. [27-29]). Hence, embryos would consume carotenoids according to their susceptibility and their response would not be 100% effective.

Here, we studied carotenoid consumption in brown trout embryos that had been experimentally exposed to *P. fluorescens* in order to test the aforementioned hypotheses. We used a new sample of offspring of the females that Wilkins et al. [24] had studied. This allowed us to link carotenoid consumptions to the life-history traits that were described in Wilkins et al. [24], and that are closely linked to two fitness-relevant traits of salmonids, namely the timing of emergence from the gravel bed and the size at emergence [30, 31]. We investigated whether carotenoids simply prevent pathogenic stress or are used by the embryo defence against the bacterial pathogen.

## 2 Methods and materials

### 2.1 Ethical note

This study complied with the relevant ethical regulations imposed by the University of Lausanne, the canton, and the country in which it was carried out. The Fishery Inspectorate of the Bern Canton granted permission for handling adults and embryos. No authorization from the cantonal veterinary office was necessary because manipulation of the adults was part of the yearly hatchery program of the Bern Canton and all experimental manipulations on embryos were performed prior to yolk sac absorption.

### 2.2 Field sampling and artificial fertilizations

Adult brown trout (37 females and 35 males) were caught at their natural spawning grounds in two connected tributaries (Kiese and Rotache) of the river Aare in Switzerland. See Stelkens et al. [32–34] for population genetic analyses of the various subpopulations in the study region. Fish were stripped at a cantonal hatchery (*Fischereistützpunkt Reutigen*) where a sample of four eggs per female was immediately frozen in liquid nitrogen for later measurements of astaxanthin, capsanthin, lutein and zeaxanthin as described in Wilkins et al. [24]). The remaining eggs were used for full-factorial *in vitro* fertilizations as in Jacob et al. [35]. Females and males were split into seven breeding blocks. We produced five breeding blocks composed of five females crossed with five males (i.e., 5 x 25 families) from the Kiese population, and two blocks of six females were crossed with five males (i.e., 2 x 30 families) from the Rotache population. After fertilizations, eggs were left undisturbed for two hours for egg hardening, then immediately transported to the laboratory where they were washed and distributed to 24-well plates for incubation in climate chambers that controlled for temperature and light [24]. These experimental families had also been subjected to a previous study that related maternally supplemented carotenoids in eggs to offspring survival under pathogen stress [24]. The present study focuses on changes in carotenoids during embryo development.

### 2.3 Experimental protocol

We sampled 24 embryos of each family and singly distributed them to individual wells of 24-well plates (Falcon, BD Biosciences, Allschwil, Switzerland) filled with 1.8 ml of autoclaved and aerated standardised water [36]. In the 24-well plates, families were distributed column-wise so each plate contained 4 embryos of 6 different families. Embryos were kept at 6.5 °C in a climate chamber with a photoperiod of 12 hours.

We exposed 12 embryos per sib group to *P. fluorescens* (PF) and sham-treated the remaining to standardised water only. PF cultures were incubated, washed and diluted as described for “PF1” in Clark et al. [37]. Embryos were exposed to the treatment with a stock solution of 0.2 ml containing 10^7^ bacterial cells/ml, yielding a final concentration of 10^6^ bacterial cells/ml.

In order to avoid a potentially selective disappearance of some phenotypes due to pathogen-induced mortality, we chose a low-virulence strain of PF [37]. We also exposed embryos to the pathogen at one of two different time points in order to spread the risk of time-point related high mortalities. This was done because the virulence of the bacterial pathogen can depend on host development stage [15]. Three breeding blocks of Kiese and one of Rotache were exposed to the treatment 20 days after fertilization (early exposure), while the remaining breeding blocks (two for Kiese and one for Rotache) were exposed 49 days after fertilization (late exposure). It turned out that early exposure led to slightly higher mortalities [24]. However, the overall mortality was so low (around 1%; [24]) that a potential mortality-induced bias in the determination of mean carotenoid consumption could be ignored.

Sixteen of these embryos were used by Wilkins et al. [24] to link various life-history traits to egg carotenoid content. Briefly, embryo survival and hatching time were noted at the day of hatching, and hatching time, hatchling length (= larval length at hatching), yolk sac volume at hatching (calculated as in Jensen et al. [38]), and larval growth (during the first 14 days after hatching) were quantified based on images. The remaining eight embryos (four PF-exposed and four sham-exposed) were available for the present study, i.e., for measuring astaxanthin, capsanthin, lutein, and zeaxanthin at different time points during embryogenesis. The embryo sampling (i.e., freezing and storing at -80°C) was performed 28 days after exposure (i.e., 48 days post fertilization) for the early exposure breeding blocks and eight days after exposure (i.e., 57 days post fertilization) for the late exposure blocks. These measurements of carotenoids were compared to the measurements on eggs of the same females recorded in Wilkins et al. [24] and allowed for an estimation of the change in embryo carotenoid content in both treatment groups.

Since this study concentrates on maternal environmental effects, only one family per female was used for carotenoid measurements. In order to avoid potentially confounding paternal effects, a random sampling of families was performed without replacement of sire identity so that the sample of each female was sired by a different male. We measured carotenoid contents from embryos of 35 maternal half sib families, which yielded 70 samples in total (i.e., two samples per family: one with embryos exposed to PF and one with controls). Absolute change in carotenoids was determined as the initial carotenoid content per egg (reported in Wilkins et al. [24]) minus the second measurement of carotenoid content per embryo. Proportional change in carotenoids was determined as the absolute change in carotenoids per embryo divided by the initial carotenoid content per egg. As reported before [24], none of the four carotenoid contents was correlated to egg weight.

### 2.4 Carotenoid extraction and quantification

Immediately before carotenoid extractions, embryos were thawed, dried, and weighed. Four embryos of each family were pooled to reach natural carotenoid concentrations that are likely to be above detection limit [24], resulting in one sample per family and treatment. Carotenoids were extracted with ethyl acetate as described in Wilkins et al. [24]. The dried extracts were protected from light and stored at -80 °C until carotenoid quantification.

Carotenoids in eggs (N = 35) had been quantified previously for Wilkins et al. [24]. Astaxanthin, capsanthin, lutein and zeaxanthin in embryos were quantified by ultrahigh-performance liquid chromatography – high resolution mass spectrometry (UHPLC-ESI-HRMS), using the same methods that had previously been used for the eggs [24]. See S1 Table for carotenoid contents in embryos and for the technical repeatability of the measurements.

### 2.5 Statistical analyses

In order to investigate whether carotenoid contents were reduced in embryos relative to eggs, we performed paired t-tests for each carotenoid we quantified. We used Spearman’s rank correlations (*rho*) to compare changes in the different types of carotenoids and to analyse how these changes are linked to initial carotenoid content of the eggs. Multivariate analyses of variance (MANOVA) were performed in order to test whether infection and time point of infection had an effect on the proportional change of the contents of the various carotenoids (after graphical verification that the assumptions of the MANOVA were not significantly violated).

When comparing changes in carotenoids to embryo phenotypes, we decided to include proportional changes in carotenoids in our statistical models rather than absolute changes, because (i) carotenoid measurements were significantly correlated, i.e., the females with a higher initial carotenoid content also showed a greater change in carotenoids and (ii) the use of proportional changes in carotenoids controls for possible confounding effects of initial amounts of carotenoids in the eggs before fertilization. Moreover, proportional change in carotenoids is an informative variable to study the role of carotenoids for stress tolerance because it reveals the extent to which embryos consumed their carotenoid reserves. The links between mean embryo survival in both environments and changes in carotenoids was analyzed with Spearman’s rank correlations. Hatching time (days), hatchling length (mm), yolk sac volume at hatching (mm^3^), and larval growth (mm) were analysed as continuous response variables in linear mixed models (LMM). In the models, treatment, and proportional change in carotenoids were included as fixed effects, and dam as random effect. When analysing response variables after hatching (i.e., hatchling length, yolk sac volume at hatching and larval growth), we included hatching time and its interaction with treatment as fixed effects in our statistical models.

The carotenoid measurements for one female yielded unexpectedly high negative changes (female ADC: mean change in astaxanthin = -43.8 nM; mean change in lutein = -52.6 nM; mean change in zeaxanthin = -65.4 nM) and would have had an extraordinary influence on the LMM (i.e., violating the model assumptions - S1 Figure shows the disproportional statistical leverage of this female for models on embryo performance). A possible explanation for this highly negative change is the low absolute carotenoid contents measured from this female, which makes the differences between the two contents more sensitive to measurement errors. Therefore, this female was removed from LMM (but not from non-parametric analyses).

To test the significance of an effect, a model including or omitting the term of interest was compared to the reference model with Akaike’s information criterion and likelihood ratio tests (LRT). All statistical tests were analysed in R v.3.1.3 [39], and mixed effect models were run with the lme4 package v.1.1.11 [40].

## 3 Results

### 3.1 Carotenoid contents

Astaxanthin, lutein, and zeaxanthin could consistently be quantified in all embryo samples (astaxanthin: 495.6 nM/egg ± 90.2 nM/embryo (means ± 95% confidence interval); lutein: 151.9 nM/egg ± 8.5 nM/embryo; zeaxanthin: 551.3 nM/egg ± 77.1 nM/embryo; S2 Fig.). No capsanthin was found above detection limit in any sample. Measurements of carotenoid contents for each maternal half sib family and repeatability estimates are presented in S1 Table.

Measurements in eggs and embryos were significantly correlated for all carotenoids and within both environments: females that had high levels of carotenoids in their eggs before fertilization also showed larger amounts of carotenoids at the late-eyed development stage of their offspring (Table 1, Fig. 1 a-c). Average astaxanthin and zeaxanthin contents in embryos were reduced relative to their average content in eggs in either treatment (astaxanthin: *t* always > 2.0, *P* always < 0.04, mean loss 231.4 nM/individual ± 51.7 nM/individual (means ± 95% confidence interval); zeaxanthin: *t* always > 3.2, *P* always < 0.002, mean loss 302.3 nM/individual ± 73.3 nM/individual). However, in both treatments, average lutein content was not significantly different between embryos and eggs (*t* always < 1.8, *P* always > 0.08, mean loss 27.5 nM/individual ± 16.0 nM/individual). The absolute change in carotenoids was correlated to initial content for all three carotenoids: greater changes were observed in the eggs of females that already had a high initial carotenoid content (Table 1, Fig. 1d-f). Figure 2 shows the relationship between pairwise changes for all three carotenoids separately for the control and the PF treatment. Astaxanthin and zeaxanthin changes were positively correlated (*rho* always > 0.38, *P* always < 0.03; Fig. 2a). The same was true for lutein and zeaxanthin (*rho* always > 0.40, *P* always < 0.02; Fig. 2b) in both treatments. For the comparison astaxanthin *vs*. lutein, a significant correlation was only found for the PF treatment (*rho* = 0.46, *P* = 0.006; Fig. 2c) but not for the control (*rho* = 0.30, *P* = 0.08; Fig. 2c). Time point of infection and of sampling did not significantly affect changes in carotenoids, neither by itself nor in interaction with treatment (Table 2). Accordingly, time point of infection and sampling were not included in any further statistical models.

**Fig 1.**
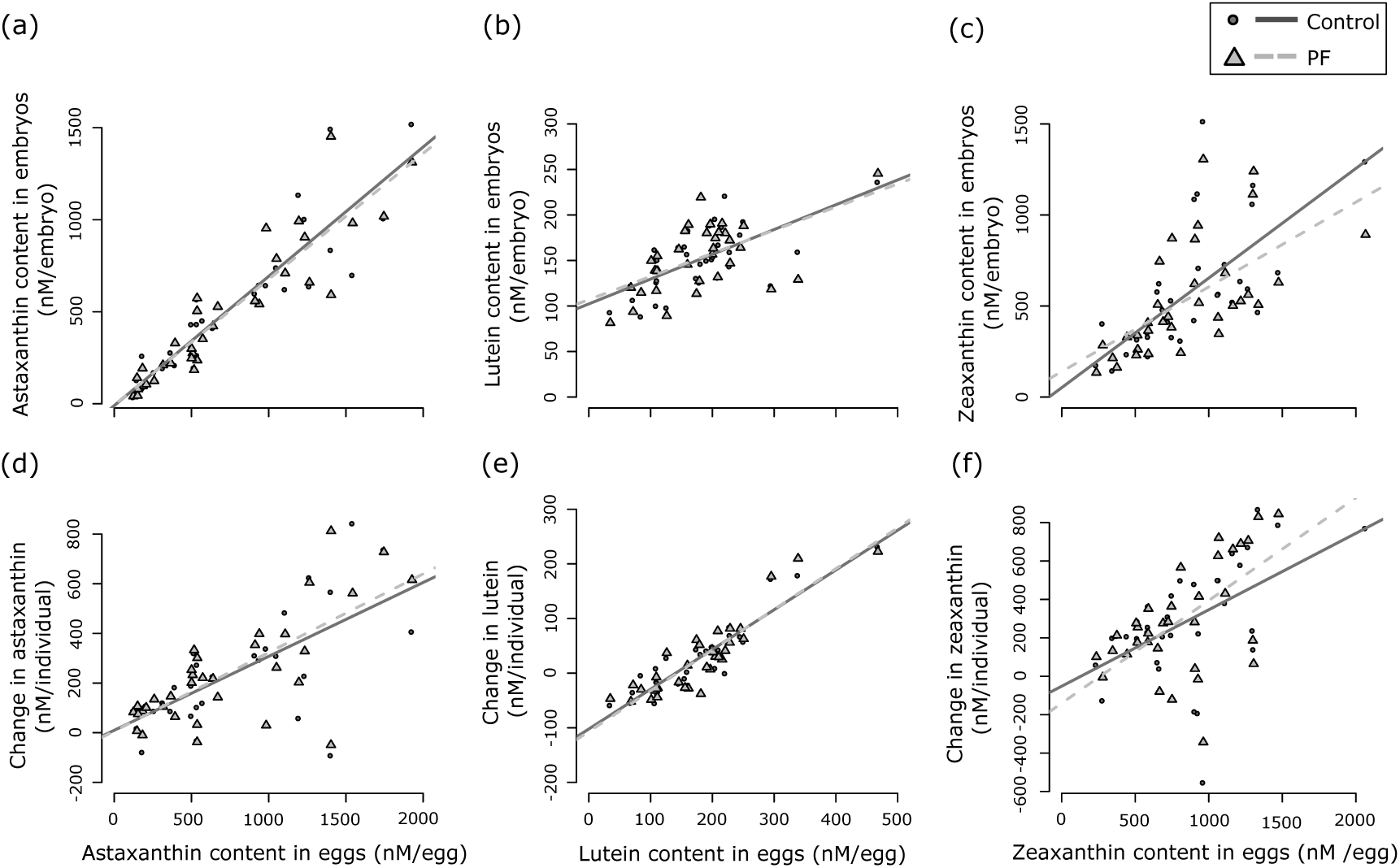
The relationship between carotenoid content in eggs and in embryos across maternal half sib families. Astaxanthin, lutein and zeaxanthin contents were measured before fertilization (“content in eggs”; data from Wilkins et al. [24]) and at a late-eyed development stage (“content in embryos”) in sham-treated controls (circles and solid regression lines) and after exposure to *P. fluorescens* (PF; triangles and dashed regression lines). Panels a-c show the relationship between carotenoid contents at the two different time points; and panels d-f the absolute changes in carotenoids relative to carotenoid contents before fertilization. See Table 1 for non-parametric statistics (the regressions lines are shown for illustration).

**Fig 2.**
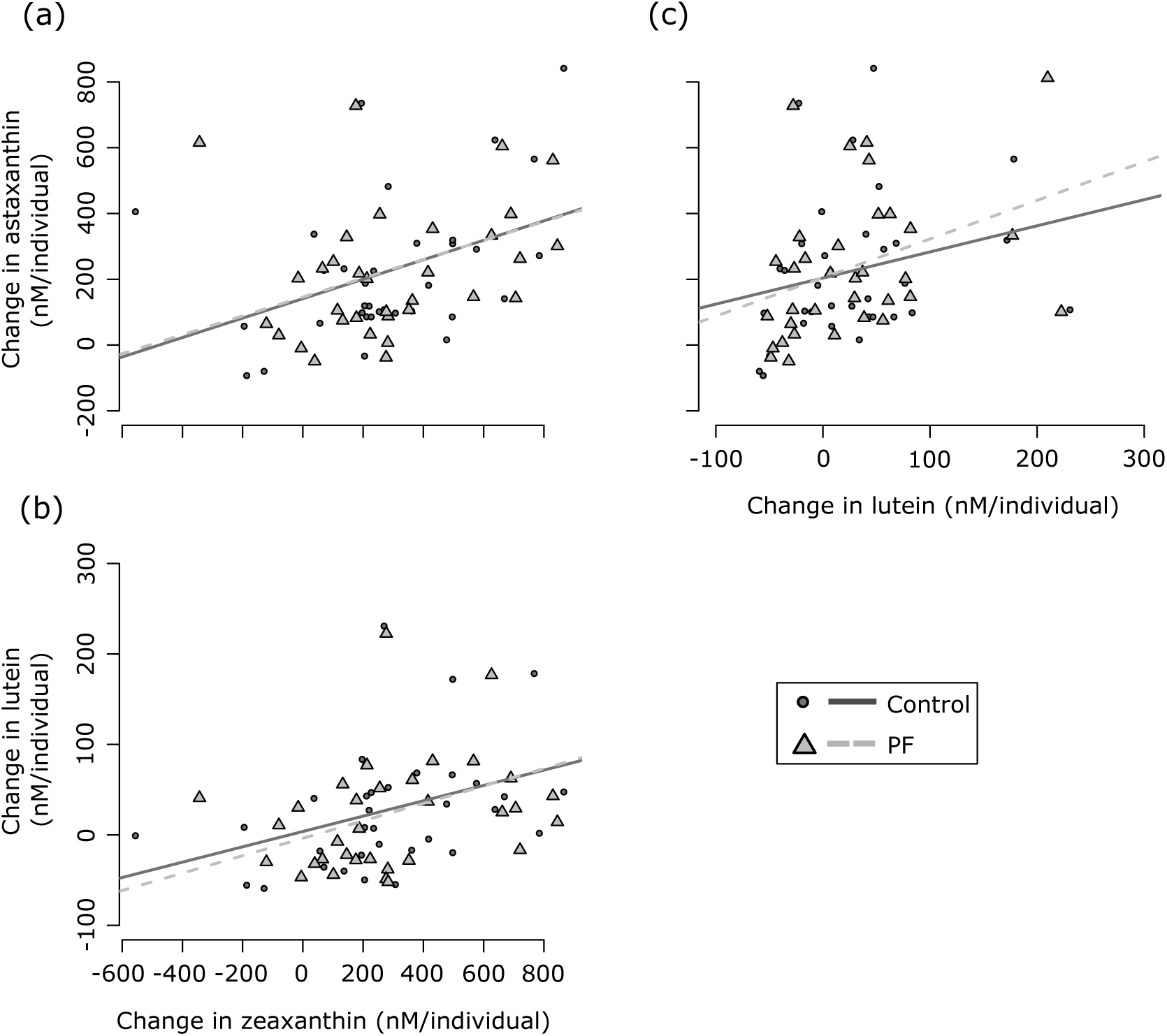
Change in carotenoid composition during embryo development. Panels represent the relationships between absolute change in (a) astaxanthin and zeaxanthin, (b) zeaxanthin and lutein, and (c) astaxanthin and lutein from fertilization to the late-eyed development stage in the sham-treated controls (circles and solid line) and in the *P. fluorescens* (PF) treated samples (triangles and dashed line). See text for non-parametric statistics. The regressions lines are shown for illustration.

**Table 1.** Spearman’s rank correlation (*rho*) between carotenoid measurements in the sham-treated controls and in *Pseudomonas fluorescens* (PF) treated embryos.

|  | Controls: content in eggs<sup>1</sup> vs. |  | PF: content in eggs<sup>1</sup> vs. |  |
| --- | --- | --- | --- | --- |
|  | content in embryos | absolute change | content in embryos | absolute change |
| Astaxanthin | 0.95*** | 0.64*** | 0.94*** | 0.64*** |
| Lutein | 0.60*** | 0.87*** | 0.51** | 0.86*** |
| Zeaxanthin | 0.78*** | 0.54** | 0.74*** | 0.54** |

<sup>1</sup>Carotenoid contents in eggs are from Wilkins et al. [24].
\*\* $P \leq 0.01$ , \*\*\* $P \leq 0.001$

**Table 2.**
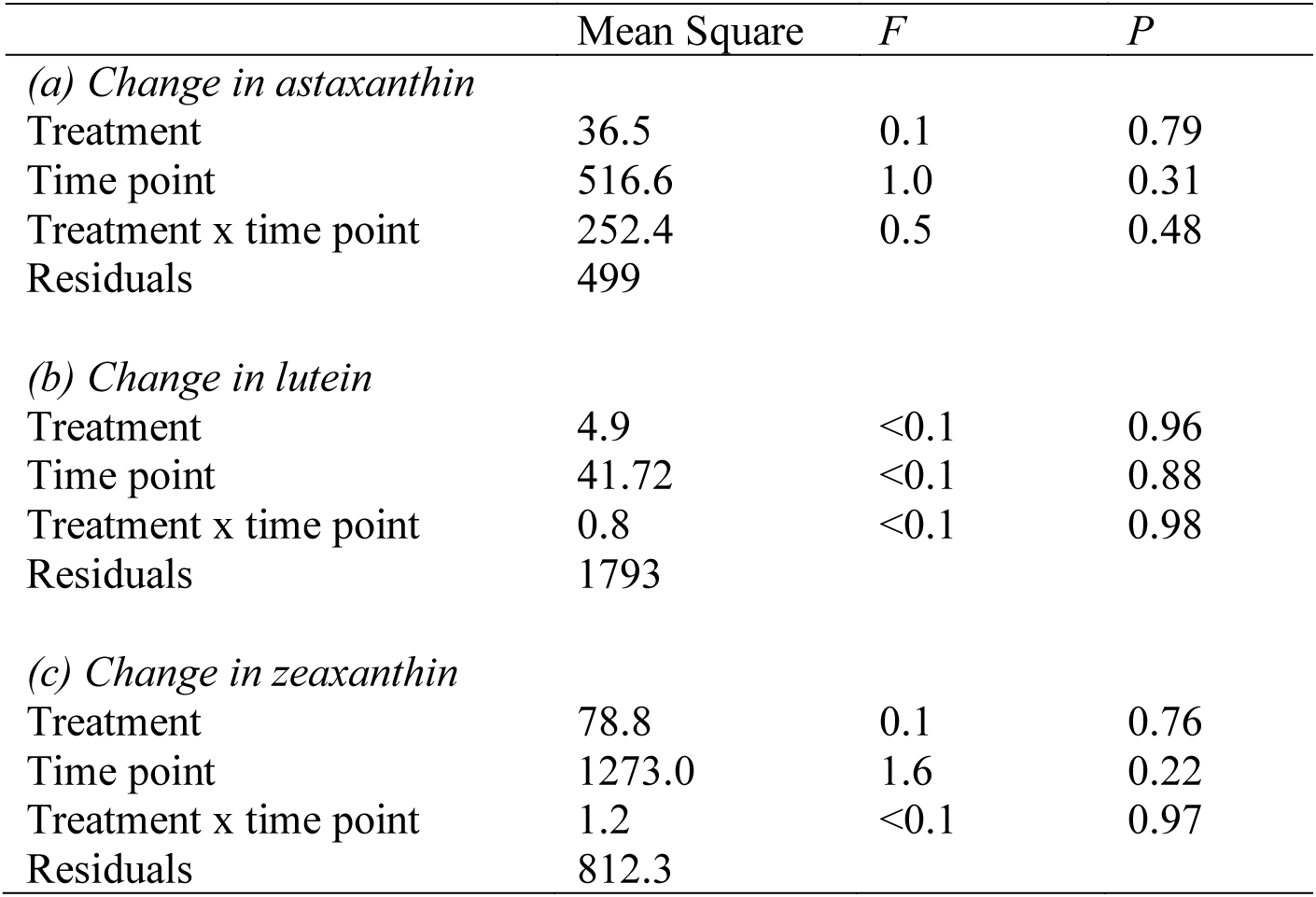
The effects of treatment, time point of infection/sampling and their interaction on (a) proportional changes in astaxanthin, (b) lutein and (c) zeaxanthin tested with a multivariate analysis of variance (MANOVA).

### 3.2 Embryo performance vs. changes in carotenoid content

Because we worked only with a subsample of the families from Wilkins et al. [24], in the present study we did not investigate the main effects of the pathogen virulence and the variance explained by dam identity on the embryo phenotypes analysed (these are reported in Wilkins et al. [24]). Here, we rather focus on whether embryo performance was linked to changes in carotenoids under a pathogen infection relative to a sham-treatment. We could not find a significant link between embryo survival and changes in astaxanthin (*rho* always between 0.23 and 0.24; *P* always > 0.17), lutein (*rho* always between -0.009 and 0.02; *P* always > 0.91), or zeaxanthin (*rho* always between 0.04 and 0.25; *P* always > 0.14) in both treatments. Proportional changes in the measured carotenoids were not significantly correlated to hatching time (Table 3a; Fig. 3a – c). No significant relationship was found between hatchling length and proportional changes in astaxanthin, lutein, or zeaxanthin under neither control nor PF exposure (Table 3b; S3a – c Fig.). Moreover, no significant links were found between hatching time and hatchling length (Table 3b; S4a Fig.). Proportional change in astaxanthin, lutein, and zeaxanthin were not significantly linked to yolk sac volume at hatching in both treatments (Table 3c; S3d – f Fig.). Hatching time was significantly correlated to yolk sac volume at hatching (Table 3c), with early hatchers tending to have larger yolk reserves than late hatchers (S4b Fig.). Hatching time also correlated to larval growth (Table 3d), with embryos that hatched later displaying faster growth than early hatchlings (S4c Fig.). However, no significant links between changes in carotenoids and larval growth were found (Table 3d; Fig. 3d – f). S5 Fig. is analogous to Fig. 3 and S3 Fig. but presents the results with the outlier female that had to be excluded from the LMM.

**Fig 3.**
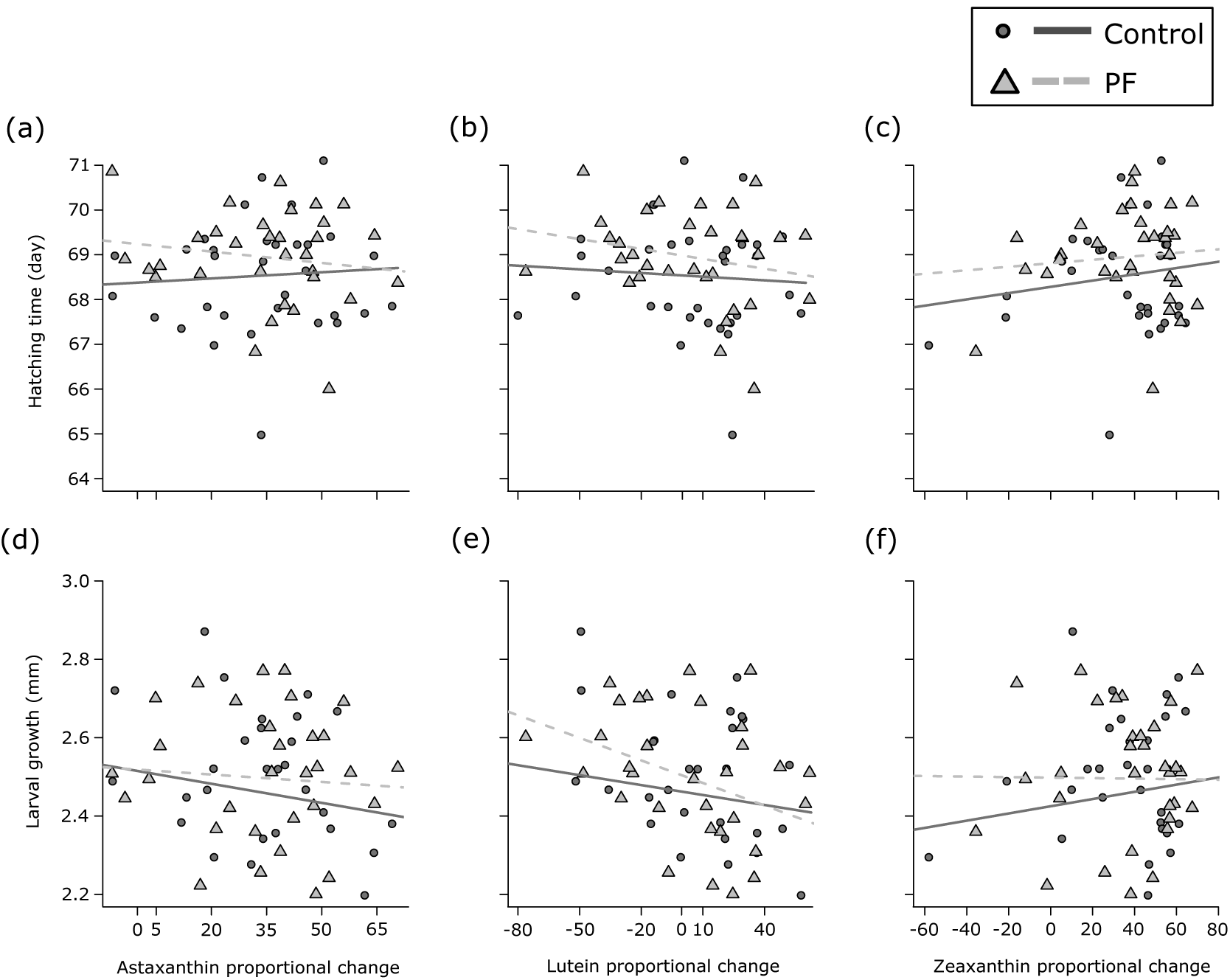
Proportional changes in carotenoid content relative to early fitness-related traits. Embryo hatching time (a – c) and larval growth (d – f) are shown for change in astaxanthin, lutein, and zeaxanthin. Changes in carotenoid contents are given for sham-treated controls (circles and solid lines) and PF treated samples (triangles and dashed lines). See Table 3 for statistics.

**Table 3.** The effects of treatment, proportional change in carotenoid contents on (a) hatching time, (b) hatchling length, (c) yolk sac volume at hatching, and (d) larval growth.

|  |  | Change in astaxanthin |  |  | Change in lutein |  |  | Change in zeaxanthin |  |  |
| --- | --- | --- | --- | --- | --- | --- | --- | --- | --- | --- |
| Model | Effect tested | AIC | $X^2$ | $P$ | AIC | $X^2$ | $P$ | AIC | $X^2$ | $P$ |
| (a) Hatching time |  |  |  |  |  |  |  |  |  |  |
| $t + c + d$ | | 1521 | | | 1519 | | | 1521 | | |
| $t + d$ | c | 1519 | 0.2 | 0.63 | 1520 | 2.4 | 0.12 | 1520 | 0.2 | 0.65 |
| $t + c + t \times a + d$ | t x c | 1523 | 0.1 | 0.76 | 1521 | 0.4 | 0.52 | 1523 | <0.1 | 0.91 |
| (b) Hatchling length |  |  |  |  |  |  |  |  |  |  |
| $ht + t + c + d$ | | 306 | | | 307 | | | 307 | | |
| $ht + t + d$ | c | 305 | 0.7 | 0.39 | 305 | <0.1 | 0.93 | 305 | <0.1 | 0.93 |
| $t + c + d$ | ht | 306 | 2.1 | 0.15 | 307 | 2.0 | 0.16 | 307 | 2.0 | 0.16 |
| $ht + t + c + t \times a + d$ | t x c | 308 | <0.1 | 0.93 | 309 | 0.1 | 0.73 | 309 | 0.3 | 0.60 |
| $ht + t + c + t \times ht + d$ | t x ht | 308 | 0.3 | 0.57 | 308 | 0.4 | 0.52 | 308 | 0.4 | 0.51 |
| (c) Yolk sac volume |  |  |  |  |  |  |  |  |  |  |
| $ht + t + c + d$ | | 3014 | | | 3016 | | | 3016 | | |
| $ht + t + d$ | c | 3014 | 2.2 | 0.14 | 3014 | <0.1 | 0.90 | 3014 | 0.7 | 0.41 |
| $t + c + d$ | ht | 3027 | 14.6 | <0.001 | 3029 | 14.2 | <0.001 | 3028 | 14.3 | <0.001 |
| $ht + t + c + t \times a + d$ | t x c | 3014 | 2.4 | 0.11 | 3016 | 2.6 | 0.10 | 3018 | <0.1 | 0.85 |
| $ht + t + c + t \times ht + d$ | t x ht | 3014 | 2.1 | 0.15 | 3017 | 1.6 | 0.20 | 3016 | 1.8 | 0.17 |
| (d) Larval growth |  |  |  |  |  |  |  |  |  |  |
| $ht + t + c + d$ | | 554 | | | 554 | | | 555 | | |
| $ht + t + d$ | c | 553 | 1.3 | 0.25 | 553 | 1.9 | 0.17 | 553 | 0.3 | 0.60 |
| $t + c + d$ | ht | 580 | 27.9 | <0.001 | 578 | 26.6 | <0.001 | 583 | 29.4 | <0.001 |
| $ht + t + c + t \times a + d$ | t x c | 556 | <0.1 | 0.88 | 555 | 0.6 | 0.42 | 557 | <0.1 | 0.99 |
| $ht + t + c + t \times ht + d$ | t x ht | 556 | 0.1 | 0.74 | 556 | 0.1 | 0.74 | 557 | 0.2 | 0.68 |

Fixed effects: t, treatment; ht, hatching time; c, proportional change in carotenoid (i.e., either astaxanthin, lutein or zeaxanthin). Random effect: d, dam.

Separate models were tested for individual carotenoids. For hatchling length, yolk sac volume at hatching and larval growth (panels b – d), models also account for the effect of hatching time. Effects were tested by comparing a model lacking or including the effect of interest to the reference model (in italics) with likelihood ratio tests. Significant effects are highlighted in bold. See Wilkins et al. [24] for the effects of treatment and dam on embryo traits.

## 4 Discussion

Studies on host-pathogen interactions often suffer from the problem that pathogen-induced mortality can lead to the selective disappearance of some phenotypes and genotypes in the samples [23]. Here, we successfully avoided this problem by using a pathogen strain that turned out to induce virulence but only very low mortality (on average about 1%, [24]). Our measurements of carotenoid consumption are therefore not confounded by non-random mortality. We found that carotenoid consumption was not significantly linked to resistance to this pathogen. Such a non-significant result can have one of the four potential explanations: (i) It reveals that defence against this pathogen does not include significant consumption of carotenoids, (ii) it is a product of large measurement error in carotenoid quantification and/or offspring phenotypes, (iii) it is based on poor measures of pathogen resistance, and (iv) it is a consequence of lack of statistical power due to insufficient sample size. In the following we argue that the first is the most parsimonious explanation of our findings.

With regards to carotenoid quantifications: We found the carotenoid contents at late embryonic stage measured here to be highly correlated to previous measurements on eggs from the same females [24]. This builds confidence in our quantification methods. Moreover, carotenoid measurements from PF-and sham-exposed embryos from the same family were highly correlated within the present study. This builds further confidence in our quantifications because carotenoids were independently extracted and quantified not only per female but also per treatment. We therefore conclude that our methods allowed for great repeatability of carotenoid quantifications.

We argue that the life-history traits that we investigated in this study are useful indicators of pathogen resistance because they are linked to the timing of emergence from the gravel and to the size at emergence. Both have been shown to be fitness-relevant in salmonids. Brown trout depend on feeding territories [41], and larvae that emerge early from the gravel bed after yolk sac consumption are more likely to establish such a territory and to outcompete late-emerging competitors [30]. Larvae that emerge larger from the gravel bed have better swimming ability and are superior competitors that can, for example, better evade predators [31]. Regarding measurement errors in these early phenotypes: Salmonid embryos have been previously used in various contexts and proved to be sensitive indicators of environmental stress. For example, not only exposure to pathogens triggers changes in various life-history parameters [13, 15] but even the sterilized odour of a pathogen infection can induce precocious hatching within a few hours [42, 43]. Other types of environmental stressors also induced significant changes in phenotypes, often at surprising low concentrations. For example, the toxicity of 17α-ethinylestradiol (EE2) could be verified in a single exposure to only 2 pg [14, 44], while previous studies had concentrated on higher doses [45, 46]. Singly-raised embryos were repeatedly used to quantify the components of phenotypic variance [e.g. 12, 15], and even within-family variation on a single genomic region could be shown to affect phenotypes under different environmental conditions [e.g. 47]. Therefore, singly reared embryos in fully controlled laboratory environments are sensitive indicators of environmental changes, and we are confident that our protocols would have allowed the detection of changes in phenotypes in response to changes in carotenoids.

With regards to sample size: We used the females and exposure protocols of Wilkins et al. [24], who quantified different aspects of embryo development in response to this pathogen. While Wilkins et al. [24] quantified carotenoids in pools of 4 eggs per female (N=35), we used pools of 4 embryos per female and treatment (i.e., a total of 70 measurements). Moreover, Wilkins et al. [24] found egg carotenoid contents (one measurement per maternal sib group) to be linked to overall offspring performance and resistance to PF. Since we linked the same observations of offspring performance to the newly determined changes in carotenoid contents (two measurements per maternal sib group), we argue that the statistical power of the present study is comparable to the one of Wilkins et al. [24] to detect correlations between virulence measures and changes in carotenoids.

Comparing the results of Wilkins et al. [24] with the present study suggests that carotenoids are useful for preventing a pathogen stress (i.e., they seem to be important at the first line of defense by reducing susceptibility) but are not significantly consumed during the immune response to the infection. We observed that carotenoids were lost over time in both treatment groups. However, we did not find significant effects of the pathogen or of the sampling time point (9 days difference) on carotenoid loss. Moreover, it remained unclear whether changes in carotenoids are only due to consumption. A loss in carotenoids can also be a consequence of degradation (when carotenoids are decayed by abiotic factors and lose their typical chemical properties) or transformation (when one carotenoid is metabolized one into another one). Carotenoids can be degraded by oxidation [48]. *In vivo* oxidation is, for example, caused by heat shock, exposure to light, or the interaction and stabilization of free radical species [48]. For some sib groups, we found that the content of carotenoids was higher in late embryonic stages than in eggs and average lutein contents did not significantly decrease throughout ontogeny, suggesting carotenoid transformations played a role during embryo development. Indeed, several carotenoids can be metabolites of other carotenoids [49]. For example, astaxanthin, lutein, and zeaxanthin can be metabolites of each other in several taxa, including fish [49–51].

The recent study by Wilkins et al. [23] found loss in carotenoids to be significantly linked to embryo mortality under increased organic pollution. The contrast between their results and ours suggests that, with regard to the role that carotenoids play during embryo development, organic pollution affects embryos differently than a single-strain pathogen infection. While a single-strain pathogen infection may largely be an immune challenge for developing embryos, organic pollution is a change in the microecology that supports symbiotic microbial communities. Increased microbial growth can directly induce virulence and/or negatively change water quality, such as a reduction in oxygen concentrations [35, 52]. Therefore, high concentrations of organic pollution typically induce significant mortality in salmonid embryos[23, 35, 52].

Similar to our study, Tyndale et al. [19] investigated in the Chinook salmon (*Oncorhynchus tshawytscha*) the loss of carotenoids during embryonic development and their role for embryo survival. The authors did not experimentally add an environmental stressor, but the mortality rates they observed suggest the presence of such a stressor. In accordance to our results, Tyndale et al. [19] found that astaxanthin decreased during development, but that its loss was not linked to embryo survival. The role of carotenoids for tolerance to environmental stress may therefore be stress-specific.

In conclusion, we tested for a link between carotenoid consumption and pathogen resistance using experimental protocols and a sample size that were sufficient to successfully establish links between initial egg carotenoid contents and pathogen resistance [24]. We found no effect of a pathogen infection on consumption of carotenoids, i.e., the infection did not induce a higher loss of carotenoids. Although carotenoids are linked to the primarily susceptibility to the pathogen we tested, they are not significantly consumed during immune response.

## 5 Acknowledgements

We would like to thank L. Benaroyo, I. Castro, P. Christe, G. Glauser, M. Hobil, K. Mobley, S. Moro, D. Nusbaumer, C. Primmer, A. Uppal, A. Vallat, and D. Zeugin for assistance in the field and/or discussion, and C. Récapet and the other two anonymous reviewers for comments on the manuscript. We also thank the staff of the Fisheries Inspectorate Berne, especially U. Gutmann and B. Bracher from the *Fischereistützpunkt Reutigen* for catching and taking care of the adult fish, and C. Küng for permissions. This study complied with the relevant ethical regulations imposed by the University of Lausanne, the canton, and the country in which it was carried out.

## Supporting information

**S1 Table.** Summary of the average concentrations (nM) of the measured carotenoids in embryo samples per maternal half sib family and treatment with associated standard deviations (SD) and coefficients of variation (CV). Data represent technical replicates (two independent runs of the same sample at different times during ultrahigh-performance liquid chromatography – high resolution mass spectrometry). The first three letters of the sample name identify the dam, the last three numbers identify the sire, “-C-” stands for sham-treated control, and “-PF-“ stands for samples exposed to PF.

| Sample | Lutein |  |  | Astaxanthin |  |  | Zeaxanthin |  |  |
| --- | --- | --- | --- | --- | --- | --- | --- | --- | --- |
|  | Average (nM) | SD (nM) | CV (%) | Average (nM) | SD (nM) | CV (%) | Average (nM) | SD (nM) | CV (%) |
| ACW-C-106 | 2.6 | 0.2 | 8 | 23.9 | 2.8 | 12 | 17.4 | 0.8 | 4 |
| ACW-PF-106 | 2.2 | 0.1 | 5 | 23.2 | 0.8 | 4 | 13.9 | 1.6 | 12 |
| ACX-C-107 | 2.4 | 0.3 | 11 | 10.3 | 1.9 | 19 | 10.0 | 0.5 | 5 |
| ACX-PF-107 | 2.9 | 0.3 | 9 | 15.3 | 0.7 | 5 | 11.9 | 0.9 | 8 |
| ACY-C-112 | 1.6 | 0.2 | 13 | 7.2 | 0.6 | 9 | 11.4 | 1.4 | 12 |
| ACY-PF-112 | 1.4 | 0.1 | 9 | 5.6 | 0.9 | 15 | 8.3 | 0.9 | 11 |
| ACZ-C-109 | 2.6 | 0.2 | 8 | 13.4 | 0.8 | 6 | 20.7 | 2.0 | 10 |
| ACZ-PF-109 | 2.1 | 0.2 | 12 | 9.5 | 1.2 | 13 | 14.3 | 1.8 | 13 |
| ADA-C-111 | 1.7 | 0.1 | 5 | 16.1 | 0.7 | 4 | 9.3 | 0.4 | 5 |
| ADA-PF-111 | 1.5 | 0.1 | 7 | 14.5 | 1.1 | 8 | 8.1 | 0.9 | 11 |
| ADB-C-113 | 2.6 | 0.3 | 13 | 9.6 | 1.0 | 10 | 11.7 | 1.8 | 15 |
| ADB-PF-113 | 2.4 | 0.2 | 7 | 8.9 | 1.1 | 12 | 10.9 | 0.2 | 2 |
| ADC-C-110 | 1.5 | 0.2 | 12 | 4.2 | 0.9 | 21 | 6.5 | 1.2 | 19 |
| ADC-PF-110 | 1.3 | 0.2 | 12 | 3.1 | 0.5 | 17 | 4.6 | 0.8 | 18 |
| ADD-C-116 | 1.6 | 0.0 | 3 | 3.1 | 0.2 | 6 | 3.8 | 0.5 | 14 |
| ADD-PF-116 | 1.9 | 0.1 | 5 | 3.4 | 0.1 | 2 | 5.3 | 0.1 | 2 |
| ADF-C-116 | 1.42 | 0.03 | 2 | 4.2 | 0.5 | 12 | 8.7 | 0.9 | 10 |
| ADF-PF-116 | 1.8 | 0.2 | 11 | 5.3 | 0.1 | 1 | 13.9 | 3.1 | 22 |
| ADG-C-114 | 2.0 | 0.3 | 14 | 10.0 | 1.3 | 13 | 3.7 | 0.3 | 8 |
| ADG-PF-114 | 2.0 | 0.2 | 9 | 11.4 | 0.8 | 7 | 4.2 | 0.7 | 16 |
| ACP-101-C-1 | 2.1 | 0.1 | 2 | 2.7 | 0.5 | 19 | 8.5 | 1.1 | 14 |
| ACP-101-PF-1 | 1.8 | 0.1 | 4 | 2.0 | 0.3 | 15 | 6.1 | 1.2 | 19 |
| ACV-104-C-1 | 2.0 | 0.1 | 4 | 1.4 | 0.1 | 5 | 6.6 | 0.8 | 12 |
| ACV-104-PF-1 | 1.9 | 0.0 | 1 | 1.5 | 0.1 | 4 | 7.1 | 0.2 | 3 |
| AEK-151-C-1 | 2.5 | 0.2 | 6 | 0.6 | 0.0 | 3 | 5.7 | 0.3 | 6 |
| AEK-151-PF-1 | 2.6 | 0.1 | 3 | 0.7 | 0.0 | 7 | 6.5 | 0.8 | 13 |
| AEM-150-C-1 | 2.0 | 0.3 | 17 | 0.7 | 0.2 | 24 | 3.6 | 0.6 | 16 |
| AEM-150-PF-1 | 2.2 | 0.1 | 4 | 0.7 | 0.0 | 7 | 3.8 | 0.5 | 12 |
| AEN-147-C-1 | 2.3 | 0.2 | 8 | 0.9 | 0.1 | 17 | 2.3 | 0.3 | 11 |
| AEN-147-PF-1 | 2.8 | 0.1 | 5 | 1.3 | 0.1 | 10 | 3.4 | 0.3 | 9 |
| AEP-148-C-1 | 3.8 | 0.1 | 3 | 1.6 | 0.1 | 4 | 3.7 | 0.2 | 5 |
| AEP-148-PF-1 | 3.9 | 0.2 | 6 | 1.7 | 0.1 | 7 | 3.7 | 0.3 | 9 |
| AER-155-C-1 | 2.51 | 0.04 | 2 | 4.2 | 0.0 | 1 | 11.0 | 0.3 | 3 |
| AER-155-PF-1 | 2.3 | 0.1 | 6 | 3.8 | 0.7 | 19 | 10.1 | 1.2 | 12 |
| AES-155-C-1 | 2.1 | 0.1 | 2 | 5.0 | 0.3 | 5 | 2.6 | 0.2 | 9 |
| AES-155-PF-1 | 2.1 | 0.1 | 4 | 4.8 | 0.7 | 15 | 2.6 | 0.2 | 8 |
| AEV-153-C-1 | 2.0 | 0.1 | 4 | 6.9 | 0.1 | 1 | 2.8 | 0.2 | 8 |
| AEV-153-PF-1 | 2.48 | 0.02 | 1 | 4.0 | 0.1 | 2 | 2.1 | 0.2 | 7 |
| ACK-96-C-1 | 3.1 | 0.2 | 6 | 10.3 | 0.3 | 3 | 10.2 | 2.3 | 22 |
| ACK-96-PF-1 | 3.0 | 0.1 | 4 | 8.7 | 1.0 | 12 | 8.4 | 0.6 | 7 |

**S1 Fig.**
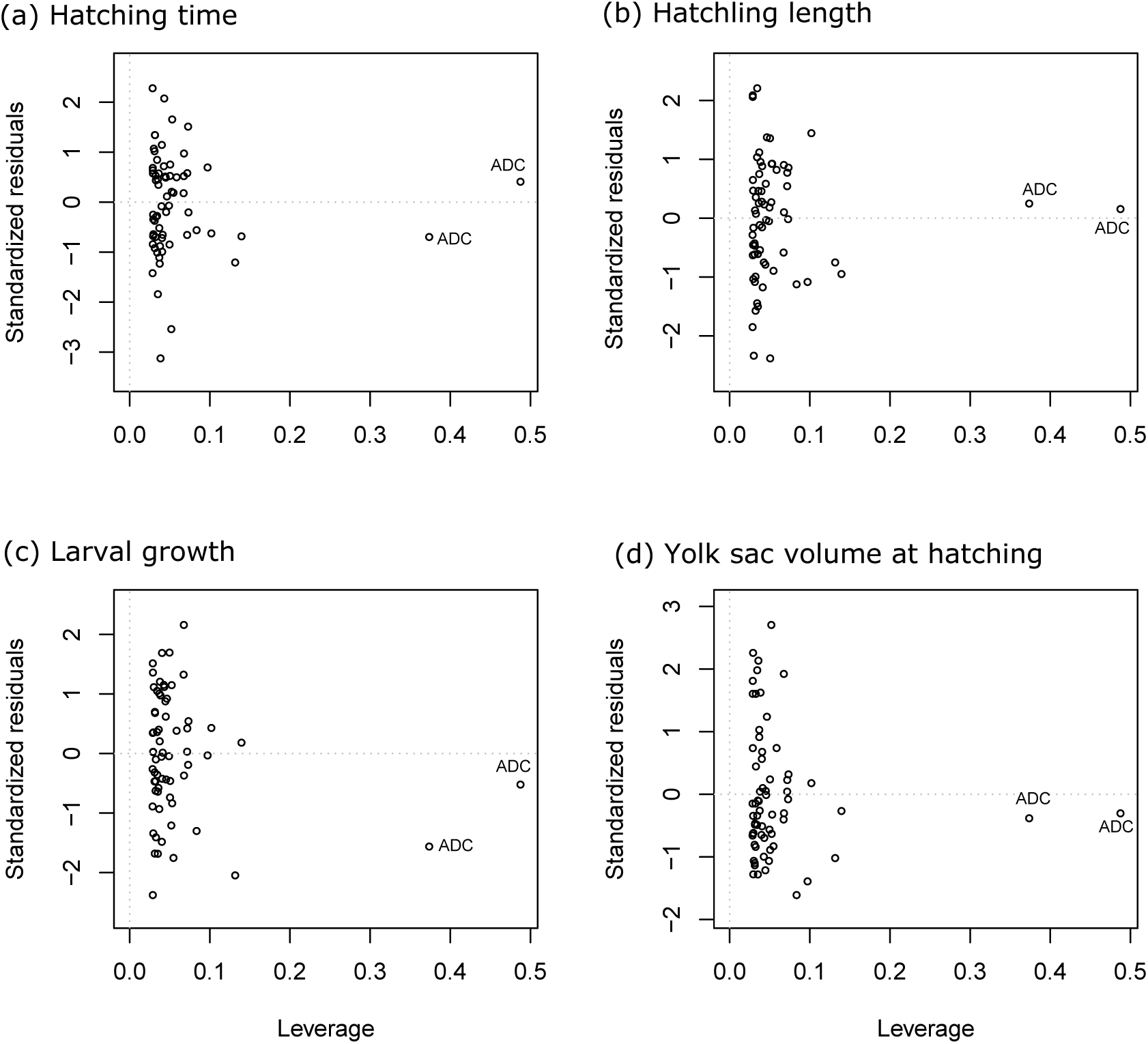
Statistical leverage relative to standardized residuals of each female in linear models on embryo performance, including the outlier female. Panels represent data of 15 models on the effects of treatment, change in lutein, and their interaction on (a) hatching time, 16 (b) hatchling length, (c) larval growth, and (d) yolk sac volume at hatching. Each female 17 represents two data points (one for PF exposure and one for sham treatment). The outlier 18 female (“ADC”) is indicated in each panel. This outlier female also presented disproportional 19 statistical leverage in models on embryo traits and change in astaxanthin.

**S2_Fig.**
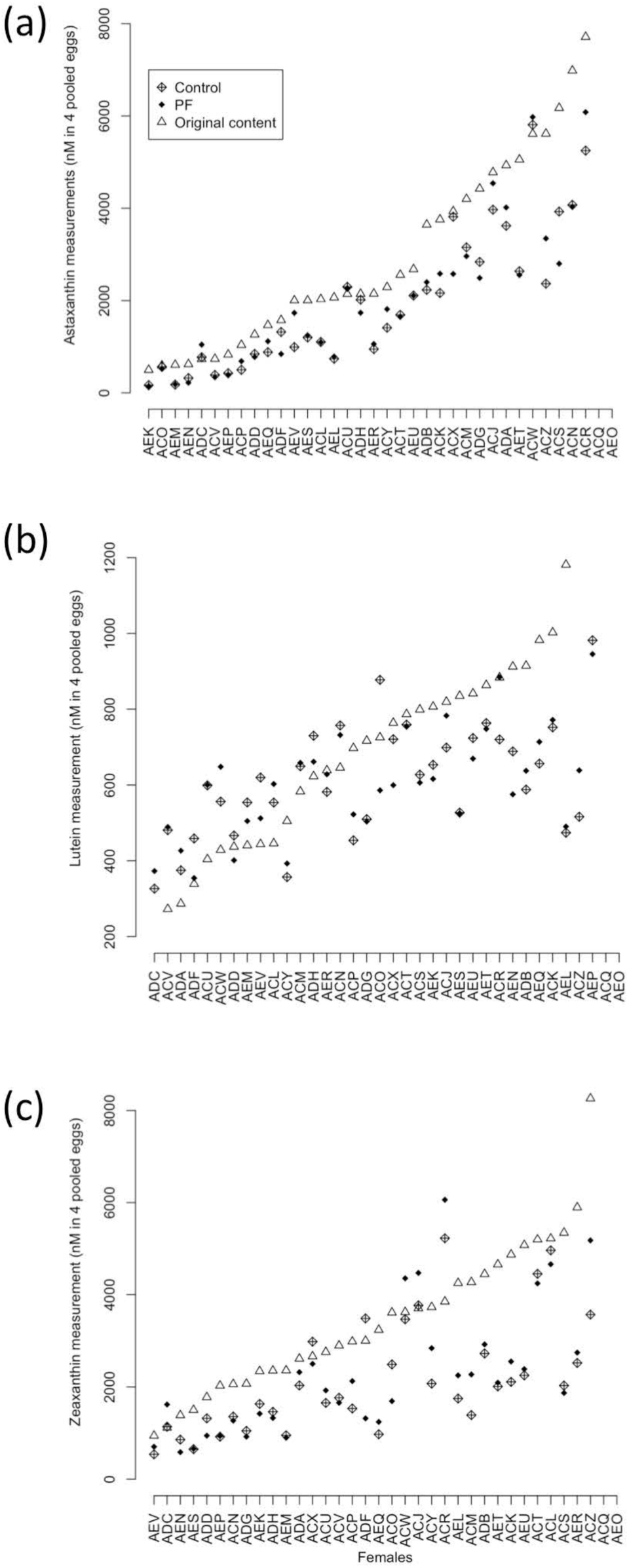
The three carotenoid measurements per female. Carotenoids were quantified in 25 eggs before fertilization (triangles), at the late-eyed development stage in the control 26 treatment (crossed diamonds) and in the PF treatment (filled diamonds) for (a) astaxanthin, 27 (b) lutein and (c) zeaxanthin. Data on the eggs come from Wilkins et al. [24].

**S3 Fig.**
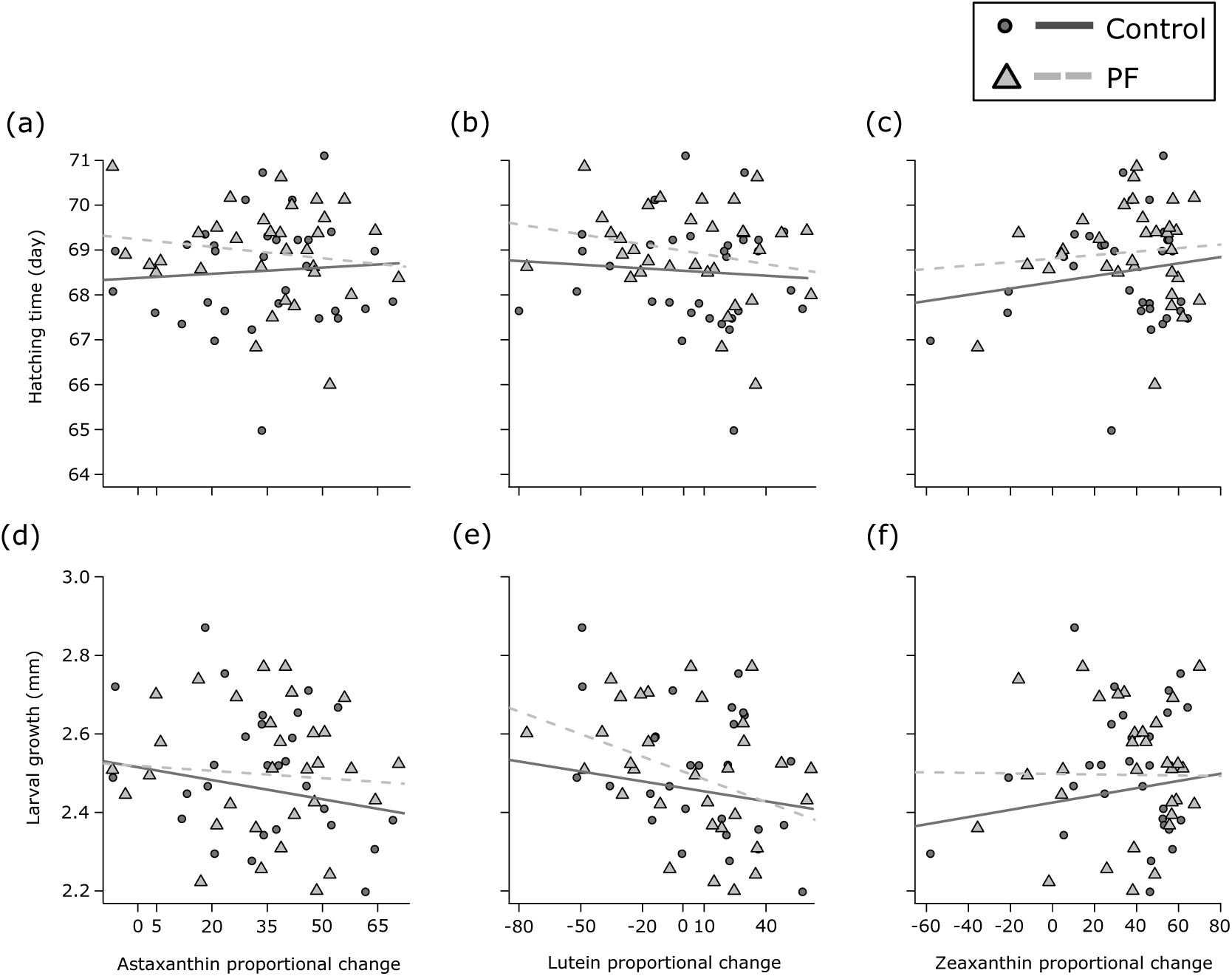
Proportional changes in carotenoid content relative to early fitness-related traits. Hatchling length (a – c) and yolk sac volume at hatching (d – f) are shown for change in astaxanthin, lutein, and zeaxanthin. Changes in carotenoid contents are given for sham-treated controls (circles and solid lines) and PF treated samples (triangles and dashed lines). See Table 3 for statistics.

**S4 Fig.**
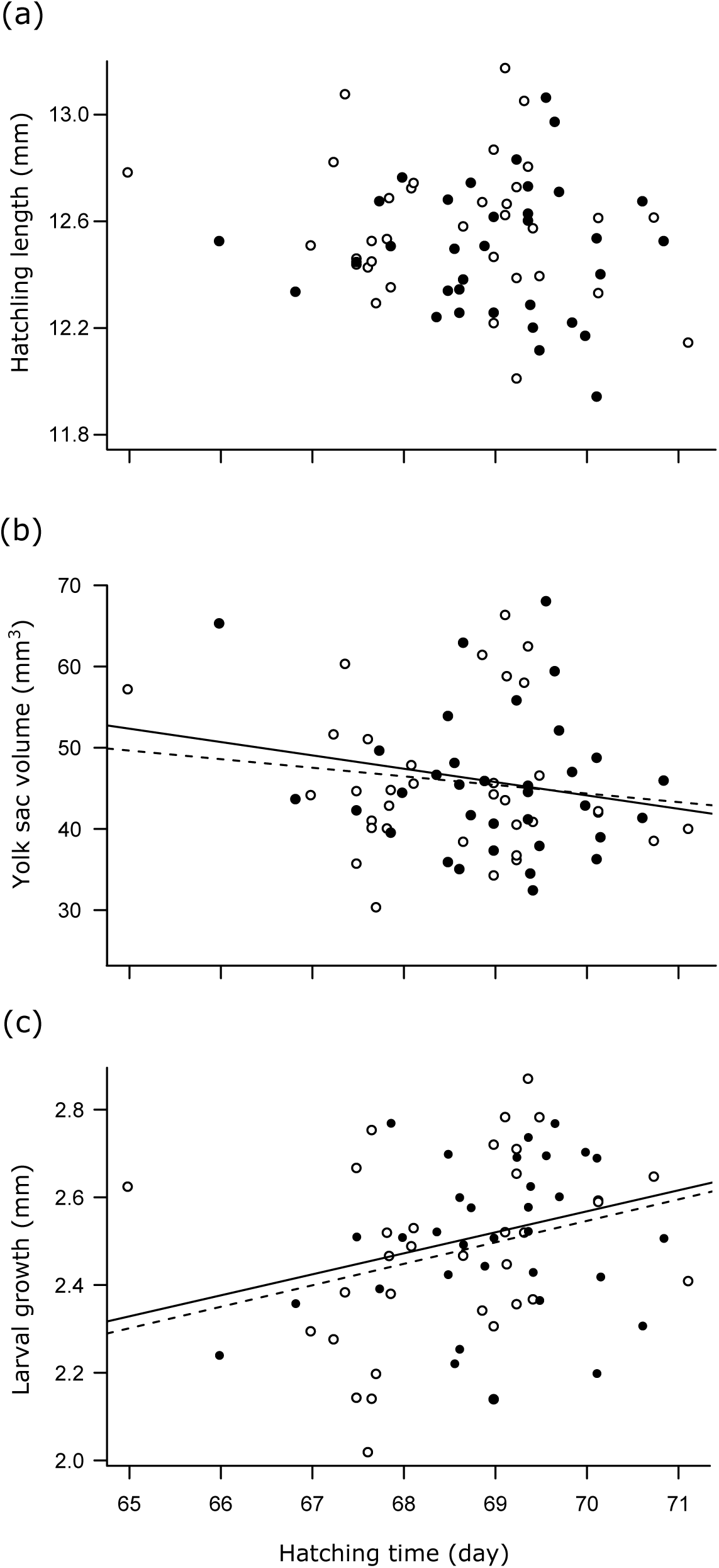
Relationship between larval traits. Means per families are shown for the control group (white circles, dotted lines) *vs*. the PF treatment (black circles, solid lines) for hatching time *vs*. (a) hatchling length, (b) yolk sac volume at hatching, and (c) larval growth. See Table 3 for statistics.

**S5 Fig.**
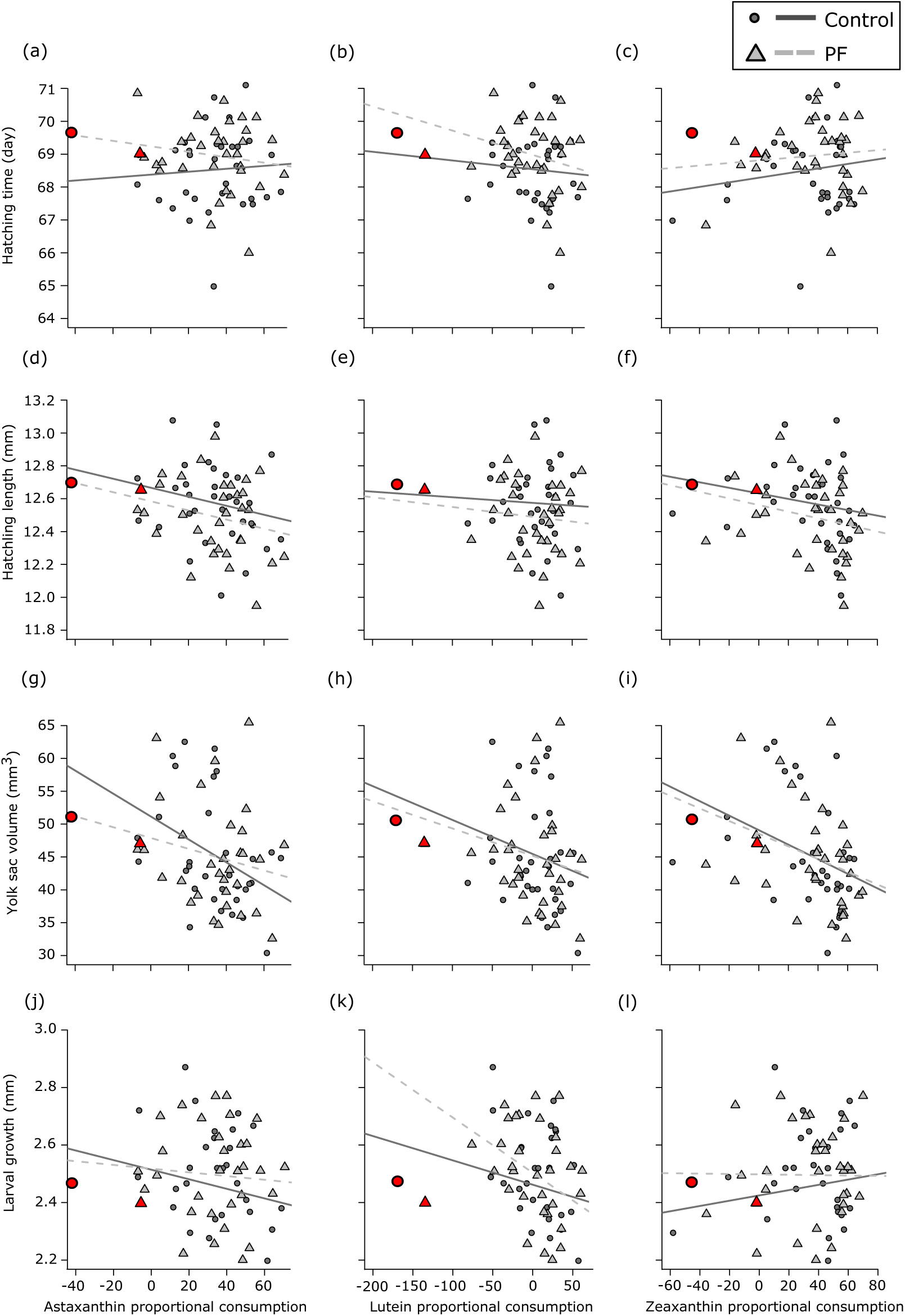
Proportional changes in carotenoid content relative to early fitness-related traits. Embryo hatching time (a – c), hatchling length (d – f), yolk sac volume at hatching (g – i), and larval growth (j – l) are shown for change in astaxanthin, lutein, and zeaxanthin. Changes in carotenoid contents are given for sham-treated controls (circles and solid lines) and PF treated samples (triangles and dashed lines). Data points of the female excluded from the analyses presented in Table 3 and illustrated in Fig. 3 and S3 are highlighted in red. Slopes correspond to the statistics in Table 3.

## References

1. Blount JD, Houston DC, Møller AP. Why egg yolk is yellow. Trends Ecol Evol. 2000;15(2):47–9. doi: 10.1016/S0169-5347(99)01774-7.

2. Ambati RR, Phang SM, Ravi S, Aswathanarayana RG. Astaxanthin: sources, extraction, stability, biological activities and its commercial applications - a review. Mar Drugs. 2014;12(1):128–52. doi: 10.3390/md12010128.

3. Olson VA, Owens IPF. Costly sexual signals: are carotenoids rare, risky or required? Trends Ecol Evol. 1998;13(12):510–4. doi: 10.1016/S0169-5347(98)01484-0.

4. Aslani BA, Ghobadi S. Studies on oxidants and antioxidants with a brief glance at their relevance to the immune system. Life Sci. 2016;146:163–73. doi: 10.1016/j.lfs.2016.01.014.

5. Palace VP, Werner J. Vitamins A and E in the maternal diet influence egg quality and early life stage development in fish: a review. Sci Mar. 2006;70:41–57.

6. Blomhoff R, Blomhoff HK. Overview of retinoid metabolism and function. J Neurobiol. 2006;66(7):606–30. doi: 10.1002/neu.20242.

7. McCallum IM, Cheng KM, March B. Carotenoid pigmentation in two strains of chinook salmon (Oncorhynchus tshawytscha) and their crosses. Aquaculture. 1987;67(3-4):291–300. doi: 10.1016/0044-8486(87)90214-6.

8. Sawanboonchun J, Roy WJ, Robertson DA, Bell JG. The impact of dietary supplementation with astaxanthin on egg quality in Atlantic cod broodstock (Gadus morhua, L.). Aquaculture. 2008;283(1):97–101. doi: 10.1016/j.aquaculture.2008.06.024.

9. Garner SR, Neff BD, Bernards MA. Dietary carotenoid levels affect carotenoid and retinoid allocation in female Chinook salmon Oncorhynchus tshawytscha. J Fish Biol. 2010;76(6):1474–90. doi: 10.1111/j.1095-8649.2010.02579.x.

10. Neff BD, Pitcher TE. Genetic quality and sexual selection: an integrated framework for good genes and compatible genes. Mol Ecol. 2005;14(1):19–38. doi: 10.1111/j.1365-294X.2004.02395.x.

11. von Siebenthal BA, Jacob A, Wedekind C. Tolerance of whitefish embryos to Pseudomonas fluorescens linked to genetic and maternal effects, and reduced by previous exposure. Fish Shellfish Immunol. 2009;26(3):531–5. doi: 10.1016/j.fsi.2009.02.008.

12. Wedekind C, Müller R, Spicher H. Potential genetic benefits of mate selection in whitefish. Journal of Evolutionary Biology. 2001;14(6):980–6. doi: 10.1046/j.1420-9101.2001.00349.x.

13. Pitcher TE, Neff BD. Genetic quality and offspring performance in Chinook salmon: implications for supportive breeding. Conservation Genetics. 2007;8(3):607–16. doi: 10.1007/s10592-006-9204-z.

14. Brazzola G, Chèvre N, Wedekind C. Additive genetic variation for tolerance to estrogen pollution in natural populations of Alpine whitefish (Coregonus sp., Salmonidae). Evol Appl. 2014;7(9):1084–93. doi: 10.1111/eva.12216.

15. Clark ES, Pompini M, Marques da Cunha L, Wedekind C. Maternal and paternal contributions to pathogen resistance dependent on development stage in a whitefish (Salmonidae). Funct Ecol. 2014;28(3):714–23. doi: 10.1111/1365-2435.12214.

16. Ahmadi MR, Bazyar AA, Safi S, Ytrestoyl T, Bjerkeng B. Effects of dietary astaxanthin supplementation on reproductive characteristics of rainbow trout (Oncorhynchus mykiss). J Appl Ichthyol. 2006;22(5):388–94. doi: 10.1111/j.1439-0426.2006.00440.x.

17. Hansen OJ, Puvanendran V, Bangera R. Broodstock diet with water and astaxanthin improve condition and egg output of brood fish and larval survival in Atlantic cod, Gadus morhua L. Aquacult Res. 2016;47(3):819–29. doi: 10.1111/are.12540.

18. Scabini V, Fernandez-Palacios H, Robaina L, Kalinowski T, Izquierdo MS. Reproductive performance of gilthead seabream (Sparus aurata L., 1758) fed two combined levels of carotenoids from paprika oleoresin and essential fatty acids. Aquac Nutr. 2011;17(3):304–12. doi: 10.1111/j.1365-2095.2010.00766.x.

19. Tyndale ST, Letcher RJ, Heath JW, Heath DD. Why are salmon eggs red? Egg carotenoids and early life survival of chinook salmon (Oncorhynchus tshawytscha). Evol Ecol Res. 2008;10(8):1187–99.

20. Anbazahan SM, Mari LSS, Yogeshwari G, Jagruthi C, Thirumurugan R, Arockiaraj J, et al. Immune response and disease resistance of carotenoids supplementation diet in Cyprinus carpio against Aeromonas hydrophila. Fish Shellfish Immunol. 2014;40(1):9–13. doi: 10.1016/j.fsi.2014.06.011.

21. Kolluru GR, Grether GF, South SH, Dunlop E, Cardinali A, Liu L, et al. The effects of carotenoid and food availability on resistance to a naturally occurring parasite (Gyrodactylus turnbulli) in guppies (Poecilia reticulata). Biol J Linn Soc. 2006;89(2):301–9. doi: 10.1111/j.1095-8312.2006.00675.x.

22. Brown AC, Cahn MD, Choi S, Clotfelter ED. Dietary carotenoids and bacterial infection in wild and domestic convict cichlids (Amatitlania spp.). Environmental Biology of Fishes. 2016;99(4):439–49. doi: 10.1007/s10641-016-0485-x.

23. Wilkins LG, Marques da Cunha L, Glauser G, Vallat A, Wedekind C. Environmental stress linked to consumption of maternally derived carotenoids in brown trout embryos (*Salmo trutta*). Ecol Evol. 2017;7(14):5082–93. doi: 10.1002/ece3.3076.

24. Wilkins LG, Marques da Cunha L, Menin L, Ortiz D, Vocat-Mottier V, Hobil M, et al. Maternal allocation of carotenoids increases tolerance to bacterial infection in brown trout. Oecologia. 2017;185(3):351–63. doi: 10.1007/s00442-017-3952-y.

25. Wilkins LGE, Rogivue A, Schütz F, Fumagalli L, Wedekind C. Increased diversity of egg-associated bacteria on brown trout (*Salmo trutta*) at elevated temperatures. Sci Rep. 2015;5. doi: 10.1038/srep17084.

26. Einum S, Fleming IA. Maternal effects of egg size in brown trout (*Salmo trutta*): norms of reaction to environmental quality. Proc R Soc Lond B Biol Sci. 1999;266(1433):2095–100. doi: 10.1098/rspb.1999.0893.

27. Løvoll M, Kilvik T, Boshra H, Bøgwald J, Sunyer JO, Dalmo RA. Maternal transfer of complement components C3-1, C3-3, C3-4, C4, C5, C7, Bf, and Df to offspring in rainbow trout (*Oncorhynchus mykiss*). Immunogenetics. 2006;58(2-3):168–79. doi: 10.1007/s00251-006-0096-3.

28. Tateno H, Yamaguchi T, Ogawa T, Muramoto K, Watanabe T, Kamiya H, et al. Immunohistochemical localization of rhamnose-binding lectins in the steelhead trout (*Oncorhynchus mykiss*). Dev Comp Immunol. 2002;26(6):543–50. doi: 10.1016/S0145-305x(02)00007-1.

29. Yousif AN, Albright LJ, Evelyn TPT. Purification and characterization of a galactose-specific lectin from the eggs of coho salmon *Oncorhynchus kisutch* and its interaction with bacterial fish pathogens. Dis Aquat Organ. 1994;20(2):127–36. doi: 10.3354/dao020127.

30. Skoglund H, Einum S, Forseth T, Barlaup BT. The penalty for arriving late in emerging salmonid juveniles: differences between species correspond to their interspecific competitive ability. Funct Ecol. 2012;26(1):104–11.

31. Einum S, Fleming IA. Selection against late emergence and small offspring in Atlantic salmon (*Salmo salar*). Evolution. 2000;54(2):628–39.

32. Stelkens RB, Jaffuel G, Escher M, Wedekind C. Genetic and phenotypic population divergence on a microgeographic scale in brown trout. Mol Ecol. 2012;21(12):2896–915. doi: doi: 10.1111/j.1365-294X.2012.05581.x.

33. Stelkens RB, Pompini M, Wedekind C. Testing for local adaptation in brown trout using reciprocal transplants. BMC Evolutionary Biology. 2012;12(1):247. doi: 10.1186/1471-2148-12-247.

34. Stelkens RB, Pompini M, Wedekind C. Testing the effects of genetic crossing distance on embryo survival within a metapopulation of brown trout (*Salmo trutta*). Conservation Genetics. 2014;15(2):375–86. doi: 10.1007/s10592-013-0545-0.

35. Jacob A, Evanno G, von Siebenthal BA, Grossen C, Wedekind C. Effects of different mating scenarios on embryo viability in brown trout. Mol Ecol. 2010;19(23):5296–307. doi: 10.1111/j.1365-294X.2010.04884.x.

36. OECD. Guideline for testing of chemicals 203 (fish acute toxicity test). Paris, France: OECD Publishing; 1992.

37. Clark ES, Wilkins LGE, Wedekind C. MHC class I expression dependent on bacterial infection and parental factors in whitefish embryos (Salmonidae). Mol Ecol. 2013;22(20):5256–69. doi: 10.1111/mec.12457.

38. Jensen LF, Hansen MM, Pertoldi C, Holdensgaard G, Mensberg KLD, Loeschcke V. Local adaptation in brown trout early life-history traits: implications for climate change adaptability. Proc R Soc Lond B Biol Sci. 2008;275(1653):2859–68. doi: 10.1098/rspb.2008.0870.

39. R Development Core Team. R: A language and environment for statistical computing. Vienna, Austria: R Foundation for Statistical Computing; 2015.

40. Bates D, Mächler M, Bolker BM, Walker SC. Fitting linear mixed-effects models using lme4. J Stat Softw. 2015;67(1):1–48. doi: 10.18637/jss.v067.i01.

41. B. J, N. J. Ecology of Atlantic salmon and brown trout. New York, USA: Springer; 2011.

42. Pompini M, Clark ES, Wedekind C. Pathogen-induced hatching and population-specific life-history response to waterborne cues in brown trout (*Salmo trutta*). Behav Ecol Sociobiol. 2013;67(4):649–56. doi: 10.1007/s00265-013-1484-y.

43. Wedekind C. Induced hatching to avoid infectious egg disease in whitefish. Current Biology. 2002;12(1):69–71. doi: 10.1016/S0960-9822(01)00627-3.

44. Marques da Cunha L, Uppal A, Seddon E, Nusbaumer D, Vermeirssen E, Wedekind C. Toxicity of 2 pg ethynylestradiol in brown trout embryos (*Salmo trutta*). bioRxiv. 2017:161570. doi: 10.1101/161570.

45. Aris AZ, Shamsuddin AS, Praveena SM. Occurrence of 17alpha-ethynylestradiol (EE2) in the environment and effect on exposed biota: a review. Environment International. 2014;69:104–19. doi: 10.1016/j.envint.2014.04.011.

46. Caldwell DJ, Mastrocco F, Anderson PD, Länge R, Sumpter JP. Predicted-no-effect concentrations for the steroid estrogens estrone, 17β-estradiol, estriol, and 17α-ethinylestradiol. Environmental Toxicology and Chemistry. 2012;31(6):1396–406. doi: 10.1002/etc.1825.

47. Wedekind C, Walker M, Portmann J, Cenni B, Müller R, Binz T. MHC-linked susceptibility to a bacterial infection, but no MHC-linked cryptic female choice in whitefish. Journal of Evolutionary Biology. 2004;17(1):11–8. doi: 10.1046/j.1420-9101.2004.00669.x.

48. Boon CS, McClements DJ, Weiss J, Decker EA. Factors influencing the chemical stability of carotenoids in foods. Crit Rev Food Sci Nutr. 2010;50(6):515–32. doi: 10.1080/10408390802565889.

49. Matsuno T. Aquatic animal carotenoids. Fish Sci. 2001;67(5):771–83. doi: 10.1046/j.1444-2906.2001.00323.x.

50. Yuangsoi B, Jintasataporn O, Tabthipwon P, Kamel C. Utilization of carotenoids in fancy carp (*Cyprinus carpio*): astaxanthin, lutein and carotene. World Appl Sci J. 2010;11(5):590–8.

51. Schiedt K, Leuenberger FJ, Vecchi M, Glinz E. Absorption, retention and metabolic transformations of carotenoids in rainbow trout, salmon and chicken. Pure Appl Chem. 1985;57(5):685–92. doi: 10.1351/pac198557050685.

52. Wedekind C, Gessner M, Vazquez F, Maerki M, Steiner D. Elevated resource availability sufficient to turn opportunistic into virulent fish pathogens. Ecology. 2010;91(5):1251–6. doi: 10.1890/09-1067.1.

